# Fetal Dehydroepiandrosterone from Hair Samples at Birth Predicts Language Development

**DOI:** 10.1101/2025.06.24.661252

**Authors:** Michaela Reimann-Ayiköz, Jasmin Preiß, Eva Reisenberger, Cristina Florea, Monika Angerer, Manuel Schabus, Dietmar Roehm, Gesa Schaadt, Claudia Männel

**Author notes:** These authors contributed equally (Gesa Schaadt, Claudia Männel). **Correspondence to:** Michaela Reimann-Ayiköz, Department of Linguistics, University of Salzburg, Erzabt-Klotz-Str. 1, 5020 Salzburg, Austria.

## Abstract

**Objective:** Sex hormones testosterone and estradiol have been related to children’s language development. Expanding the focus on dehydroepiandrosterone (DHEA), which has not yet been considered as a biological marker of language ability, may provide novel insights, as emerging evidence suggests that fetal DHEA plays a critical role in the organization of the neonatal brain, potentially shaping later language development. The present study investigated whether fetal DHEA, compared to fetal testosterone, is associated with infant language development.

**Design and methods:** DHEA and testosterone concentrations were measured in newborn hair strands (*n* = 63) collected two weeks after birth, capturing fetal long-term hormone secretions during the third trimester of pregnancy. At six months of age, children’s language abilities were assessed using the German version of the Bayley Scales of Infant Development. **Results.** Linear regression analysis revealed fetal DHEA levels to be significantly associated with language abilities at six months of age, with lower DHEA levels corresponding to higher language scores. Control analyses assessing general cognitive abilities showed no association of fetal DHEA levels with infant cognitive function. Testosterone levels were not associated with language.

**Conclusions:** The current study identifies fetal DHEA levels extracted from newborn hair samples as a potential biological factor influencing infant language development.

## 1 INTRODUCTION

Sex hormones have an essential impact on the hemispheric organization and functional lateralization of the developing brain, thus affecting language processing (Auyeung et al., 2013; Friederici et al., 2008; McCarthy, 2008). Previous research on the role of sex hormones in language development has focused on both prenatal and postnatal levels of testosterone and estradiol. At the prenatal stage, fetal testosterone levels extracted from amniotic fluid were linked to language comprehension in girls at four years of age, with an inverted U-shaped relationship (Finegan et al., 1992). In boys, higher perinatal testosterone concentrations from umbilical cord blood indicated smaller vocabulary at two years of age (Hollier et al., 2013) as well as an increased risk of language delay at one and three years of age (Whitehouse et al., 2012), possibly explained by higher prenatal testosterone exposure leading to atypical cerebral lateralization (Hollier et al., 2014b). The impact of postnatal sex hormone concentration on language development has received even more attention. Across sexes, early postnatal testosterone levels from saliva have been reported to negatively relate to later expressive vocabulary at 18 to 30 months of age (Kung et al., 2016) and, when measured from blood samples, to language comprehension at four years of age (Schaadt et al., 2015). For female sex hormones, evidence suggests estradiol to be positively linked to language development. In boys and girls, postnatal estradiol levels from blood samples have been positively associated with infants’ cry melody patterns and babbling skills during the first year of life (Quast et al., 2016; Wermke et al., 2014) and with language comprehension at four years of age (Schaadt et al., 2015). Notwithstanding these findings, the predictive value of prenatal and postnatal sex hormones for language abilities has not been consistently substantiated across all studies (Auyeung et al., 2009; Farrant et al., 2013). For example, Knickmeyer et al. (2005) reported that fetal testosterone drawn from amniotic fluid was not related to language abilities at four years of age. In sum, initial evidence partly suggests that testosterone and estradiol contribute to early language development. Yet, studies differ with respect to method and time point of hormone extraction and deliver inconsistent results.

Expanding the focus beyond these sex hormones to include dehydroepiandrosterone (DHEA) may open new perspectives on the hormonal basis of language development. Despite the fact that DHEA, together with its sulfate ester DHEA-S (jointly referred to as “DHEA(S)”, unless otherwise specified), is the most abundant steroid hormone in the fetus (Parker, 1999; Stárka et al., 2015), and is implicated in fetal brain development, it has rarely been addressed in research on child neurodevelopment (Greaves et al., 2019; Kamin and Kertes, 2017). DHEA(S) is mainly produced by the fetal adrenal glands beginning in the eighth to tenth week of gestation (Ishimoto and Jaffe, 2011; Mesiano and Jaffe, 1997). DHEA(S) production increases throughout the second and third trimesters, with fetal DHEA-S synthesis reaching approximately 200mg daily in this developmental period, highly exceeding maternal synthesis of this hormone during late pregnancy (Ishimoto and Jaffe, 2011; Mesiano and Jaffe, 1997; Siiteri and MacDonald, 1963; Tagawa et al., 2004). Following parturition, the structure of the adrenal cortex is dramatically remodeled: Initiated by an apoptotic process, a programmed cell death, the fetal zone atrophies within the first weeks postpartum. As the fetal zone disappears, DHEA(S) levels decline rapidly, reaching a nadir six months postnatally (Mesiano and Jaffe, 1997; Sucheston and Cannon, 1968).

In addition to its adrenal origin, DHEA can be synthesized de novo in the human brain (Zwain and Yen, 1999), with animal studies suggesting especially high levels during fetal brain development (Quinn et al., 2016). As a neuroactive steroid (Baulieu, 1998), fetal DHEA(S) has the potential to shape later development by acting on several neurotransmitter receptors, including sigma (σ_1_), NMDA, and GABA_A_ receptors (Maninger et al., 2009; Quinn et al., 2018). For example, DHEA(S) enhances brain maturation by stimulating the NMDA receptor through σ_1_ receptor agonism, which in turn triggers calcium entry into embryonic neocortical neurons that promotes neurite growth, emphasizing its role in early brain organization (Baulieu, 1998; Compagnone and Mellon, 2000; Compagnone and Mellon, 1998; Quinn et al., 2016). By reducing GABA_A_-ergic inhibition, DHEA(S) increases neuronal excitability (Maninger et al., 2009), potentially modulating mood (Genud et al., 2009), inflammatory processes (Hamidovic et al., 2024; Lee et al., 2024), and cognitive functions including memory (Jacob, 2019; McGaugh et al., 1990). Moreover, DHEA(S) exerts anti-glucocorticoid effects on the brain, thereby protecting against glucocorticoid-induced neurotoxicity in brain structure and function (Cardounel et al., 1999; Kimonides et al., 1999), a mechanism that may promote memory and learning processes (Fleshner et al., 1997).

Given its involvement in a wide range of function within the human brain (Stárka et al., 2015), there is growing research interest to link DHEA(S) to neurophysiological processes, particularly cognitive abilities, with research on adults yielding mixed findings: While some studies found DHEA to be positively related to verbal fluency (Yamada et al., 2010) and memory (Alhaj et al., 2006; Do Vale et al., 2014), other studies found no effect of DHEA(S) on cognitive abilities in adults (Carlson and Sherwin, 1999; Wolf et al., 1997a, 1997b). The inconsistencies in findings may be explained by differences in assessing endogenous versus administered DHEA(S), varying in doses and timing, alongside with variation in population characteristics, including cognitive profiles and age. Turning to early developmental stages, only one study so far has identified DHEA-S in an indirect pathway, linking placental inflammatory signals to cognitive and language outcome at 12 months through cord blood DHEA-S levels in children from low socioeconomic backgrounds, whereas DHEA-S was not directly associated with language or cognitive outcomes. While these findings provide initial insight into potential early-life mechanisms of DHEA(S) on a child’s neurocognitive development, research in this area is still in its infancy (Greaves et al., 2019; Kamin and Kertes, 2017).

The elevated levels of DHEA(S) concentration during the fetal period directs research attention to this stage of development (Bailey et al., 2024; Tegethoff et al., 2011), which can be captured using newborn hair samples as a retrospective method of fetal hormone analysis that is not influenced by circadian or situational variations (Gao et al., 2013; Gareri and Koren, 2010; Hart et al., 2023). In direct comparison, even though DHEA-S levels at birth, measured in cord blood, are around 400 times higher than those of DHEA (Troisi et al., 2003), DHEA appears to be a more effective biomarker than DHEA-S (Hart et al., 2023; Tegethoff et al., 2011). Crucially, DHEA is more responsive to short-term endocrine changes, while DHEA-S maintains more stable systemic levels and supplies active DHEA as needed (Longcope, 1995; Rosenfeld et al., 1975). Furthermore, DHEA can easily cross the blood-brain barrier, providing greater direct bioavailability to the brain, and potentially modulates central nervous system outcomes (Stárka et al., 2015). Thus, DHEA measured in hair samples seems a more reliable long-term biomarker than DHEA-S, reflecting fetal hormonal activity in the third trimester.

The current study investigated whether fetal DHEA extracted from newborn hair is associated with infants’ language development at six months of age. We hypothesized fetal DHEA to be positively related to language abilities in infants, reflecting its role as a neurosteroid involved in fetal neurocognitive development, with behavioral evidence supporting its impact on cognitive functions (for review, see Maninger et al., 2009). To control for domain-specificity, cognitive abilities are additionally measured to evaluate whether potential effects would be specific to language abilities. As previous research concentrated on testosterone in the prenatal period, we also investigated infants’ language development in relation to fetal testosterone. Based on previous evidence suggesting high testosterone levels to negatively contribute to early language development, particularly in boys (Hollier et al., 2013; Kung et al., 2016; Whitehouse et al., 2012), we expected a significant negative association between fetal testosterone and later language skills, with a more pronounced effect in males.

## 2 METHODS

### 2.1 Participants

Participants were mother-infant dyads from an ongoing longitudinal study funded by the Austrian Science Fund (FWF), which examined the effects of biological and environmental factors on infant cognitive and socio-emotional development. To this aim, pregnant women and their offspring were followed from the 34^th^ week of gestation until 12 months postpartum. Prior to participation, the experimental procedure was approved by the ethics committee of the University of Salzburg (EK-GZ: 12/2013) and mothers provided informed consent. For this study, the criteria for maternal inclusion were: (a) German as native language, (b) age of 18 years or older, (c) no pregnancy complications, except for transient findings in *n* = 3 included cases (i.e., resolved placenta previa; cervical insufficiency without further clinical relevance; hyperemesis and abnormal nuchal translucency with unremarkable follow-up NIPD result). None of the women reported smoking or drinking alcohol during pregnancy. The infant inclusion criteria were: (a) no developmental concerns in cognitive or language domains according to pediatric evaluation, (b) gestational age of at least 37 weeks at birth, (c) German as environmental language, (d) no hearing impairment at the age of six months, (e) absence of family history of language impairments (first- and second-degree). These criteria were ensured via a study-specific anamnesis questionnaire, along with questions on demographic information, such as the mother’s relationship status with the child’s father and maternal education. The demographic information and characteristics of the final sample are provided in Table 1 with gestational age at birth, infant age at hair collection, and infant biological sex being statistically considered as covariates.

**Table 1:** Participant and demographic information of final sample (*n* = 63)

| Variable |  |
| --- | --- |
| **Maternal and family characteristics** |  |
| Maternal age at delivery, mean (SD) [range] | 32.95 y (4.54) [20-42] |
| Maternal relationship status at the 34<sup>th</sup> GW (%) |  |
| Married to the child's father | 50.8 |
| In a relationship with the child's father | 44.4 |
| Single | 4.8 |
| Professional qualification of mothers (%) |  |
| Without vocational qualification | 1.6 |
| Completed vocational training | 17.5 |
| Higher vocational education | 19.0 |
| University degree (or higher) | 61.9 |
| **Infant characteristics** |  |
| Sex (% female) | 49.2 |
| Gestational age at birth in weeks, mean (SD) [range] | 39+6 (8d) [37-42] |
| Delivery mode (%) |  |
| Vaginal delivery | 61.9 |
| Operative vaginal delivery (forceps or vacuum) | 11.1 |
| C-section | 27.0 |
| Birth weight, mean (SD) [range] | 3393 g (417) [2540-4740] |
| Apgar Score, 10 min post-birth, mean (SD) [range]* | 9.97 (0.25) [8-10] |
| Age at hair collection, mean (SD) [range] | 16.54 d (3.57) [12-33] |
| Age at six months, mean (SD) [range] | 6.24 M (0.26) [5.87 – 6.97] |
| Hair mass, mean (SD) [range] | 2.38 mg (1.98) [0.2-7.5] |
| Hair length, mean (SD) [range]* | 1.96 cm (0.72) [1.0-3.5] |

*Note.* Maternal and family characteristics include maternal age, relationship status, and professional qualification. Infant characteristics comprise sex, age at birth, age at hair collection, age at the six-month BSID-III assessment, birth details, hair mass, and hair length (\* one missing value).

A total of 67 mother-infant dyads initially participated in the project. Four dyads were excluded from analysis for the following reasons: pre-term birth (*n* = 1), parental refusal to provide newborn hair sample (*n* = 1), maternal progesterone supplementation during pregnancy (*n* = 1), and smoking during pregnancy (*n* = 1). In *n* = 57 infants, at least one of the two tested sex hormones (testosterone and DHEA) were available from the newborn hair sample. As *n* = 10 DHEA levels and *n* = 16 testosterone levels were below the lower limit of quantification, these values were imputed for further analysis. As presented in Figure 1, the final participant sample included *n* = 63 (31 females).

**Figure 1.**
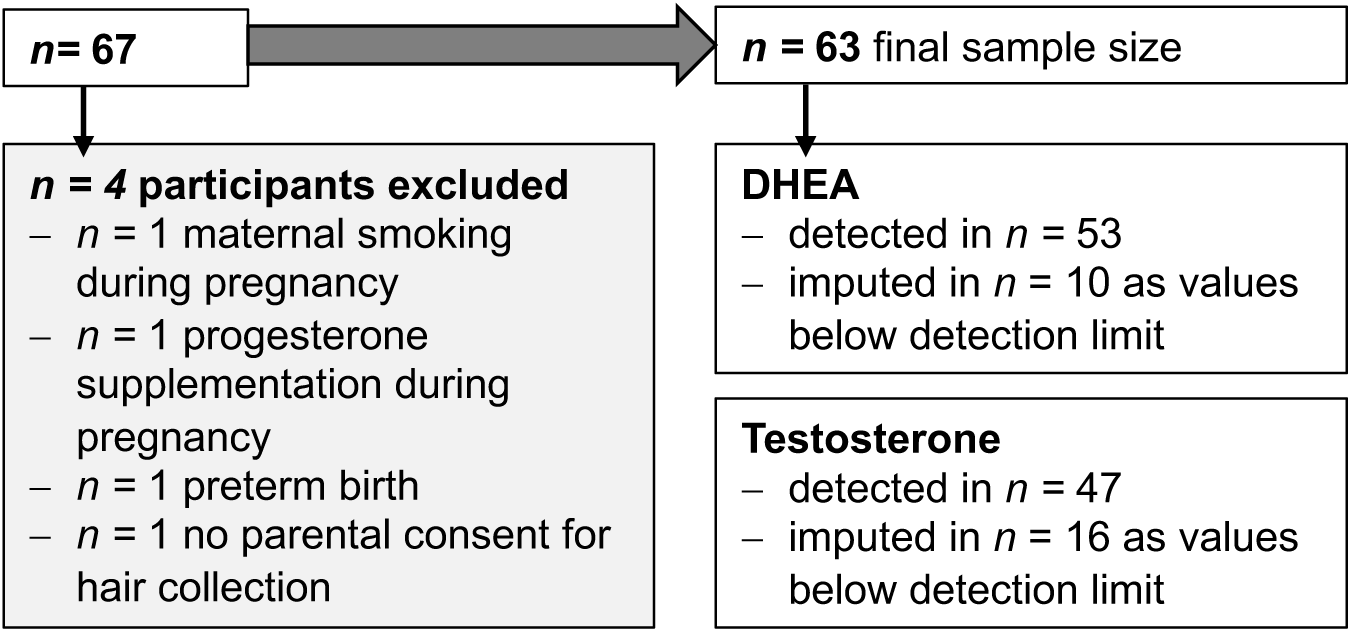
Flow chart of sample inclusion and exclusion*Note.* Following the exclusion from further analysis of four participants from the initial 67 study participants, the final sample comprised *n* = 63 participants. For DHEA, a total of *n* = 53 values were recorded, with 10 values below the detection limit addressed through imputation. For testosterone, *n = 47* cases were observed, and 16 values below the quantification limit were imputed.

As the effect of fetal DHEA on infant language abilities has not yet been tested, we estimated relevant effect sizes based on previous studies on testosterone and estradiol in order to approximate the effect sizes required for our power analysis (approximation based on two predictors, *α* = .05, *power* = 0.8). Specifically, we referred to Kung et al. (2016) for testosterone’s role in early language development measured by vocabulary at around two years (*f^2^* = 0.174, *required sample size* = 55), and Wermke et al. (2014) for estradiol’s role on infant vocal melody properties at two months of age (*f^2^* = 0.575, *required sample size* = 17), providing the basis for defining the necessary sample size (*n* = 55), which was met in the present study with *n* = 63.

### 2.2 Determination of fetal hair testosterone and DHEA

To determine fetal endogenous testosterone and DHEA levels, we measured hormone concentrations in newborn hair strands collected two weeks after birth (*M_age_* = 16.3 d, *SD* = 2.9 d). Scalp hair samples were cut at the posterior vertex, wrapped in aluminum foil, and stored in the dark. The hair sample was analyzed by the Dresden LabService GmbH at the Technische Universität Dresden, using column-switching liquid chromatography tandem mass spectrometry assay with atmospheric-pressure chemical ionization (LC-APCI-MS/MS) (Gao et al., 2013): Contrary to the procedure in adults, entire hair strands were examined in our sample, as newborn hair is in the early growth stage. Hair samples were washed with isopropanol and methanol (Carl Roth GmbH & Co. KG; Karlsruhe, Germany), incubation was used to extract testosterone and DHEA from the whole, unpowdered hair. A column switching strategy for on-line solid phase extraction (SPE) was applied for subsequent analyte detection on an AB Sciex API 5000 QTrap mass spectrometer. Stock solutions were prepared with testosterone and DHEA from Sigma-Aldrich (Hamburg, Germany), and standard solutions with testosterone-d5 and DHEA-d4 from Biocrates Life Sciences AG (Innsbruck, Austria) as deuterated internal standards. The intra-assay coefficient of variation (CV) was within an acceptable range (i.e., 9% to 12%), meeting the *Food and Drug Administration*’s acceptance limit of ≤15% CV (FDA and CDER, 2018). The lower limit of detection was 1 pg/mg for DHEA and 0.15 pg/mg for testosterone.

In contrast to hormone analyses from saliva or umbilical cord blood, the analysis of newborn hair provides a retrospective index of integrated long-term hormone secretion (Gao et al., 2013). As research on prenatal drug exposure indicates that neonate scalp hair reflects metabolic function after the 28^th^ week of gestation (Gareri and Koren, 2010), the current values can be considered as indicators of fetal hormone concentration in the third trimester of pregnancy.

### 2.3 Infant language and cognitive development

At six months of age (*M*age *=* 6.2 M, *SD* = 0.2 M), children’s neurocognitive development was assessed using the German version of the Bayley Scales of Infant Development (BSID-III) (Bayley, 2006a, 2006b; Reuner and Rosenkranz, 2014). Internal consistency, as measured by Cronbach’s alpha, was .86 in the language scale in the German norming study (*N* = 1009). The assessment was carried out by a trained female examiner in the presence of one parent. To test infants’ language skills, the language scale with the two subtests receptive and expressive communication was used (*α* = .77 and .83, respectively). The receptive communication subtest measures auditory acuity, joint attention, and responsiveness to language. The expressive communication assesses preverbal communication skills, including babbling, gesturing, joint referencing, and turn-taking. The raw scores from the subscales were first transformed into standardized scores (*M* = 10, *SD* = 3). The total score for the language scale, calculated as the sum of both standardized scores for receptive and expressive communication, was then converted to age-normed scores (*M* = 100, *SD* = 15).

To determine whether the postulated hormone-language association was domain-specific or indicative of a more general relationship with cognitive functioning, we additionally assessed cognitive abilities using the BSID-III cognitive scale. The cognitive scale for six-month-olds targets sensorimotor development and infants’ exploration and manipulation of objects and has an internal consistency of *α* = .82. Children’s raw score was transformed into a standardized, age-normed score (*M* = 100, *SD* = 15).

### 2.4 Data Analysis

Multiple linear regression analyses were used to evaluate the association between fetal DHEA and language development at six months of age, including the hormone variable as main effect as well as interaction effect with sex. In addition to the primary analysis, two control analyses were conducted: One examining cognitive development at six months of age as dependent variable, to control for whether the association with DHEA was specific to language or indicative of general cognitive abilities and the second assessing fetal testosterone as the hormone variable to examine its relationship with language development, enabling comparison with DHEA based on prior evidence of testosterone’s role in to early language outcomes (Hollier et al., 2013; Whitehouse et al., 2012). As hair length and infant age at hair sample extraction (see Table 1) were considered as potential confounders of hormone levels, we tested their associations with both hormone measures using bivariate Pearson correlations. Since neither was significantly associated with DHEA or testosterone levels (*all p >* 0.05), hair length and infant age at hair sample extraction were not included as covariates in the regression analyses (see Table S1 for detailed results).

Prior to statistical analyses, we applied multiple imputation to the hormone values below the limit of quantification for *n* = 10 participants showing DHEA values below the limit and *n* = 16 participants showing testosterone values below the limit, as outlined in the algorithm described by Herbers et al. (2021) and implemented in the R package *lnormimp* (Rupprecht, 2024). This method fits a log-normal distribution to account for the typical right-skewed pattern observed in biological data. Such a pattern was also evident in the data presented here, with significant skewness of both testosterone (skewness = 1.62, z = 4.00. p < .001, shapiro-Wilk: *W* = .81, *p < .*001) and DHEA values (skewness = 0.96, z = 2.80. p < .01, shapiro-Wilk: *W* = .93, *p <* .001). Models were estimated using multiple imputation of missing data, based on *m* = 20 imputations as recommended for missingness rates below 30% (Graham et al., 2007). Separate regression analyses were conducted on the *m* = 20 imputed datasets, and subsequently pooled using Rubin’s rules (1987) using the *miceadds* package to summarize the regression results across imputations.

Assumptions of linear regression were tested prior to model interpretation: Residuals showed homoscedasticity in all regression models, and variance inflation was consistently below 1.5, indicating no multicollinearity. Statistical outliers, defined as standardized residuals > 2.5 *SD*, were excluded in sensitivity analyses to confirm the robustness of findings (see Tables S2b, S4b). All statistical analyses were conducted using RStudio 2024.04.2+764, with a significance level of *p* = .05 (see Tables S2-S4 for detailed statistics on the multiple imputation results).

## 3 RESULTS

Concentrations of DHEA and testosterone, along with the performance in the language and cognitive scales of the BSID-III, are presented in Table 2, with no statistically significant sex differences observed in any of the variables.

**Table 2:** Fetal hormone concentrations and performance on the language and cognitive scales (*n* = 63).

| | Final sample | males | females | $p$ |
| --- | --- | --- | --- | --- |
| | ($n = 63$) | ($n = 32$) | ($n = 31$) | |
| Fetal testosterone level | 1.49 (1.86) | 1.14 (1.38) | 1.85 (2.21) | .14<sup>+</sup> |
| (pg/mg) | [0.06-8.49] | [0.06-4.97] | [0.06-8.49] |  |
| Fetal DHEA level | 17.44 (13.10) | 15.42 (13.26) | 19.53 (12.81) | .17<sup>+</sup> |
| (pg/mg) | [0.53-55.51] | [0.59-55.51] | [0.53-52.58] |  |
| BSID-III Cognitive scale | 114.10 (14.74) | 112.80 (14.75) | 115.50 (14.85) | .51<sup>+</sup> |
|  | [80-135] | [80-130] | [95-135] |  |
| BSID-III Language scale | 114.90 (11.55) | 113.30 (12.71) | 116.50 (10.17) | .28 |
|  | [85-141] | [85-141] | [94-141] |  |
| Receptive subscale raw | 9.70 (1.54) | 9.63 (1.72) | 9.77 (1.36) | .48<sup>+</sup> |
| scores, mean | [7-14] | [7-14] | [8-13] |  |
| Expressive subscale | 9.84 (1.83) | 9.69 (2.05) | 10.00 (1.59) | .35<sup>+</sup> |
| raw scores | [6-16] | [6-16] | [7-13] |  |

*Note.* For the final sample ( $n = 63$ ), the mean, standard deviation, and range for fetal testosterone and DHEA levels, as well as for the scores on the BSID-III cognitive and language scales at six months of age, are provided for each sex, with no statistically significant sex differences observed according to either the t-test or the Wilcoxon rank-sum test<sup>+</sup>. For hormone values below the quantification limit, the mean across $m = 20$ imputed datasets was used ( $n = 10$ for DHEA, $n = 16$ for testosterone).

Multiple linear regression analysis showed that the model including fetal DHEA and infant sex interacting with fetal DHEA reached significance in predicting language abilities at six months of age (*adjusted R^2^* = .13, *p <* .01, see Table S2a). Specifically, fetal DHEA was found to be negatively associated with infants’ language abilities at six months of age (*β* = -.28, *p* = .02) suggesting that the higher the fetal DHEA levels, the lower the later language abilities (see Figure 2). Note, that the interaction of fetal DHEA levels with sex was not confirmed as a significant predictor (*β* = -.23, *p* = .06) in this model, yet the statistical trend suggested a stronger association between DHEA levels and later language abilities for male infants (*β* = - .23, *p* = .06). Sensitivity analyses with statistical outliers excluded confirmed the statistically predictive value of fetal DHEA for later language abilities (see Table S2b). The control analysis, using general cognitive abilities at six months as dependent variable in the model (*adjusted R^2^* = -.01, *p* = .50, see Table S3), did not reveal any predictive value of fetal DHEA levels (*β* = .11, *p* = .39) or infant sex interacting with fetal DHEA (*β* = -.12, *p* = .36).

**Figure 2.**
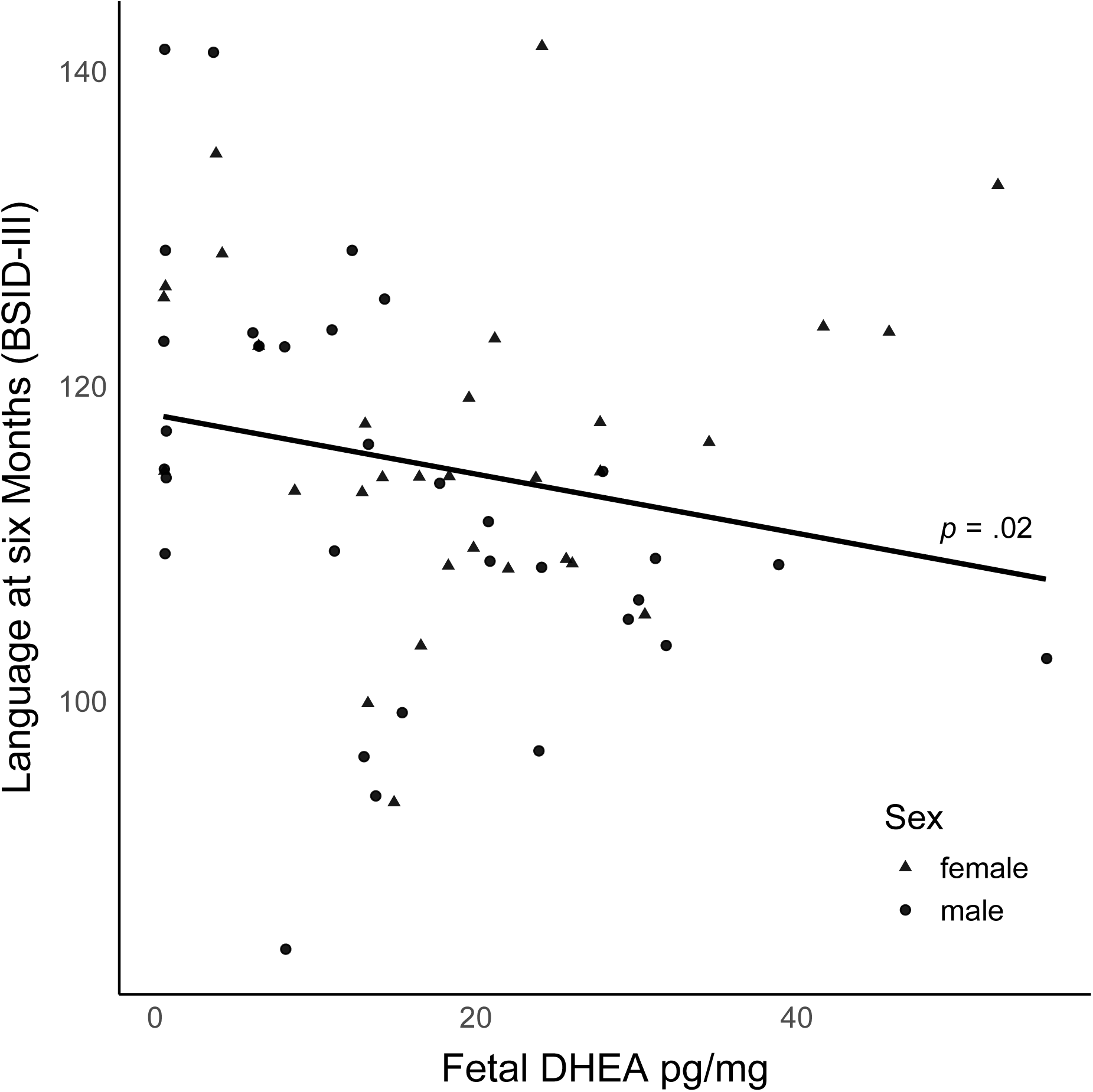
Association between fetal DHEA levels and language abilities at six months assessed using the Bayley Scales of Infant Development (BSID) III (*n* = 63). Lower language skills at six months were associated with higher levels of fetal DHEA extracted from hair at birth (*n* = 10 imputed values included, mean scores across imputed datasets shown, data jittered to prevent overlapping points).

In contrast to fetal DHEA, fetal testosterone (*β* = .16, *p* = .30, see Figure 3) was not statistically significantly associated with infants’ language abilities, nor was infant sex interacting with fetal testosterone levels (*β* = -.13, *p* = .40), as shown in multiple linear regression analysis (*adjusted R^2^* = -.01, *p* = .52, see Table S4a). Sensitivity analysis excluding outliers confirmed these results (see Table S4b).

**Figure 3.**
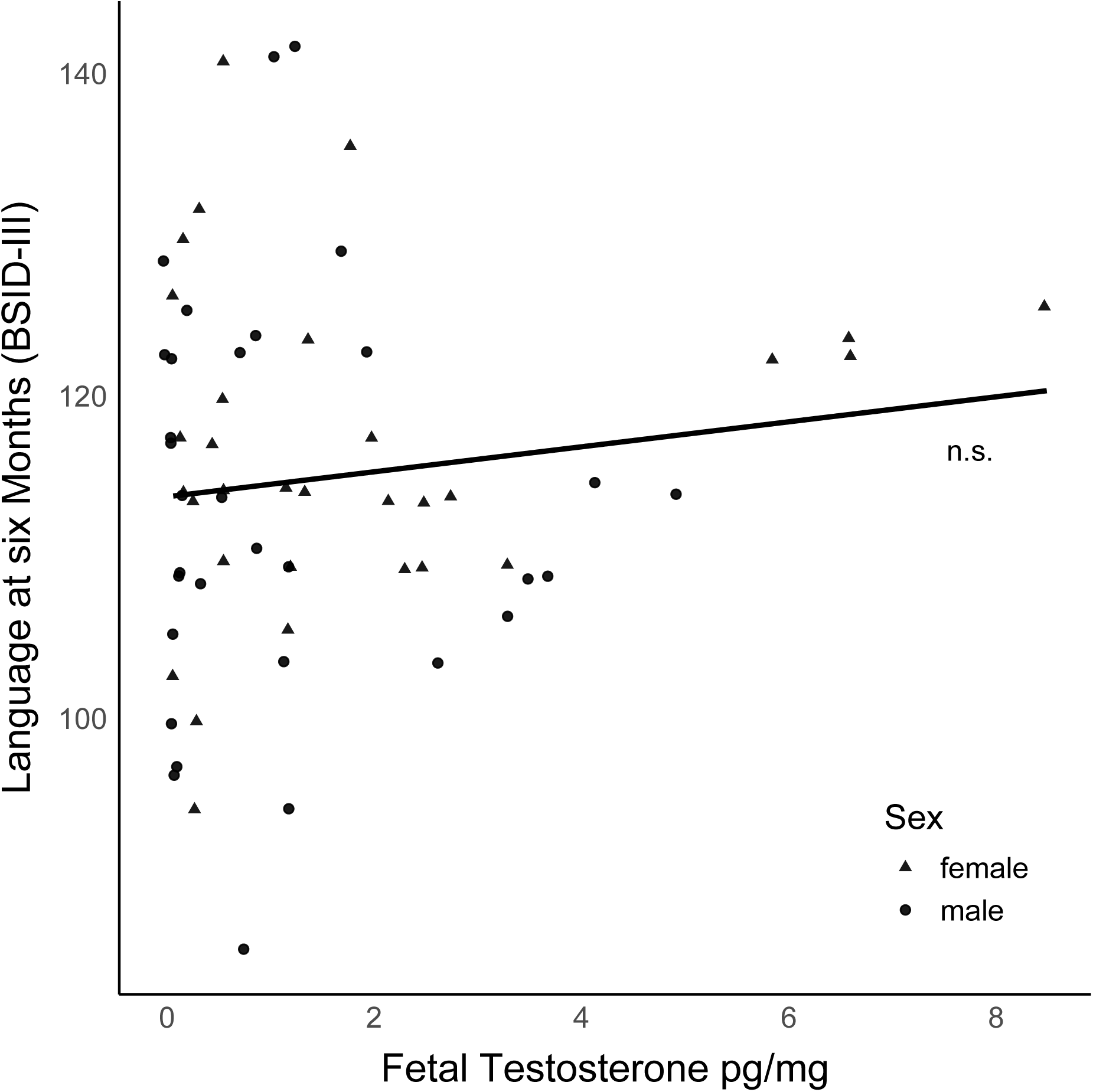
Association between fetal testosterone levels and language abilities at six months assessed using the Bayley Scales of Infant Development (BSID) III (*n* = 63). Language skills at six months were not significantly associated with fetal testosterone levels extracted from hair at birth (*n* = 16 imputed values included, mean scores across imputed datasets shown, data jittered to prevent overlapping points).

## 4 DISCUSSION

The present study examined the impact of fetal DHEA, in comparison to testosterone, on behavioral language outcome during infants’ first year of life and found that higher fetal DHEA was related to lower language outcome at six months of age. This finding suggests fetal DHEA concentration in the third trimester to serve as an early biological predictor of infant language abilities. To test for the observed effect’s language-specificity, we also evaluated general cognitive abilities, finding no significant association with DHEA. In contrast to DHEA, fetal testosterone was not associated with early language abilities. Our study not only points towards DHEA as an early biological predictor of infant language abilities, but also highlights the use of newborn hair samples for hormone extraction as a method to retrospectively infer fetal hormone concentrations during the third trimester even months before sampling is possible in newborns.

As DHEA is considered to play a crucial role in fetal brain development, particularly in processes involved in memory and learning (for review, see Maninger et al., 2009), we hypothesized fetal DHEA to be positively related to language abilities in infants, based on research in adults reporting positive associations between DHEA(S) and cognition. Contrary to the direction of this hypothesis, our results revealed a significant negative association, with higher fetal DHEA levels leading longitudinally to lower language abilities. Our findings indicate that the role of DHEA(S) in neurodevelopment may differ across developmental stages, with its function in the fetal environment likely distinct from that in later periods (Campbell, 2020). The observed negative effect of fetal DHEA levels on early language development may reflect specific mechanisms during the fetal period, possibly involving GABA_A_-receptor and anti-glucocorticoid pathways.

In the brain, DHEA(S) acts as an antagonist of the GABA_A_-receptor, which inhibits neuronal firing by increasing a CI-conductance (Maninger et al., 2009). However, in neonatal hippocampal neurons, GABA_A_ excites rather than inhibits neurons due to a reversed intracellular chloride gradient (Cherubini et al., 1991). During early development, GABA_A_, serving as the main excitatory drive, promotes neuronal growth and the formation of neuronal networks (Cherubini et al., 1991; Peerboom and Wierenga, 2021). Given that DHEA may reduce GABA_A_-driven excitation in the fetal brain, elevated DHEA levels might attenuate neuronal growth, potentially explaining the negative association with language development. However, previous studies have questioned the concept of excitatory GABA_A_ action during early development, as it has not been consistently observed in *in vivo* studies, suggesting the findings may result from methodological decisions related to brain slicing (Bregestovski and Bernard, 2012). Future studies are necessary to confirm whether GABAA acts excitatory in the fetal brain and to understand how DHEA may affect this pathway and, in turn, neurodevelopment.

In addition to the effect on GABA_A_-receptors, DHEA(S) exerts anti-glucocorticoid effects in the hippocampal brain (Cherubini et al., 1991; Peerboom and Wierenga, 2021). Regarding glucocorticoids in late gestation, the physiological increase in cortisol plays a crucial role in promoting the maturation of organs, including the brain, thereby supporting fetal neurodevelopment, while also preparing the fetus for birth (Stoye et al., 2020; Trejo et al., 2000). Indeed, maternal cortisol levels in late gestation have been found to be positively associated with vocabulary development in early childhood (Mumm et al., 2023) and cognitive functioning in middle childhood (Davis et al., 2017). Given that DHEA can counteract glucocorticoid effects, higher fetal DHEA levels may limit cortisol-driven brain development, offering an explanation for our findings linking higher DHEA in the third trimester to lower language abilities. By contrast, lower DHEA levels may enhance cortisol-driven development in the fetus.

Mechanistically, DHEA may regulate fetal cortisol exposure through its effects on the enzyme 11ß-hydroxysteroid dehydrogenase (11ß-HSD) type 2 (Maninger et al., 2009). This enzyme acts as a partial barrier in the feto-placental unit, thereby regulating fetal exposure to maternal cortisol. In late gestation, with cortisol supporting brain maturation, fetal cortisol exposure rises through decreased 11ß-HSD-2 activity, heightening fetal sensitivity to maternal cortisol levels (Howland et al., 2017, Marciniak, 2011; Murphy et al., 2003). Building on this regulatory mechanism, DHEA acts on 11ß-HSD-2 expression (Balazs et al., 2008), such that higher DHEA levels increase 11ß-HSD-2 activity, thereby limiting fetal cortisol transfer, which may adversely influence brain maturation during late gestation. With respect to our findings, elevated fetal DHEA in the third trimester may then contribute to lower language abilities through upregulation of 11ß-HSD-2 activity, thereby antagonizing cortisol-driven brain maturation, ultimately affecting early language development.

Regarding potential sex-difference, our data suggest boys may be more susceptible to the effects of DHEA on language development (*p* = .06). This pattern may reflect a potential sex-specific modulatory effect of DHEA on fetal cortisol, with cord blood cortisol correlating with placental 11ß-HSD-2 only in boys (Yu et al., 2022). Accordingly, DHEA’s modulation of 11ß-HSD-2, which in turn partially controls cortisol exposure, may play a more prominent role in male fetuses than in females, though the underlying pathways remain to be fully understood. Therefore, future studies that simultaneously assess fetal cortisol and DHEA, while considering possible sex-specific effects, will provide deeper insights into both hormones’ joint effects on neurodevelopment (Bailey et al., 2024).

An intriguing observation in our study is that fetal DHEA is specifically associated with language abilities, but not with general cognitive abilities. Most previous studies, conducted in adults (e. g. Carlson and Sherwin, 1999; Do Vale et al., 2014), have focused on cognition, challenging a direct comparison with our study. To our knowledge, so far only one study has examined both language and cognitive outcomes in infants at 12 months, reporting DHEA-S at birth being indirectly related to both domains (Lee et al., 2024). However, a direct comparison with our results is limited due to methodological differences of previous studies, including a low socioeconomic-status cohort, measuring DHEA-S from cord blood instead of fetal DHEA, and capturing cognitive abilities at a later age. Extending previous research, our study offers new insights by suggesting that also DHEA may contribute to early language development.

In contrast to DHEA, fetal testosterone levels have previously been investigated with respect to later language development employing different extraction methods which have, however, yielded contradictory results. Studies using amniotic fluid from the second trimester reported a negative association with vocabulary at age two years (Lutchmaya et al., 2001), and, in girls only, an inverted U-shaped relationship with sentence comprehension at age four years (Finegan et al., 1992). In follow-up studies with the same cohort as Lutchmaya, Baron-Cohen, and Ragatt’s (2001), the association between fetal testosterone and language abilities was no longer detectable at the age of four years (Auyeung et al., 2009; Knickmeyer et al., 2005). Moreover, studies examining perinatal testosterone levels from umbilical cord blood consistently reported a negative association with language abilities in boys, but not in girls (Hollier et al., 2013; Whitehouse et al., 2012). For example, Whitehouse and colleagues (2012) discussed higher perinatal testosterone levels as a potential risk factor for language delay in boys, and as a potential protective factor in girls. In sum, most of the studies assessing the statistical predictive value of prenatal or perinatal testosterone levels on later language outcomes found sex-specific differences, not only in terms of an association per se but also the direction of this association (Finegan et al., 1992; Whitehouse et al., 2012). In our study, however, we first, did not find an association between fetal testosterone and postnatal language skills and second, sex did not significantly add to explaining later language outcomes. Our findings may be influenced by the time window covered by hormone extraction from hair: We assessed testosterone levels during the last trimester, whereas previous studies indicating predictive effects of testosterone measured either in early pregnancy (Lutchmaya et al., 2001), at birth (Whitehouse et al., 2012), or in the early postnatal stage (Schaadt et al., 2015). This may explain the absence of predictive value of testosterone within our sample. The sex-specific predictive value of testosterone may depend on the stage of development, as testosterone levels in boys rise prenatally from gestational weeks eight to 24, peaking at 16 weeks, and rise again postnatally during the so-called mini-puberty from weeks four to 12 postpartum (Hines, 2020). Therefore, these time windows may be particularly valuable for assessing the sex-specific effects of testosterone on later development in boys and girls.

In assessing fetal hormone exposure, there is currently no methodological gold standard and different measures may lead to inconsistent results, making comparisons across studies challenging (Hollier et al., 2014a; van de Beek et al., 2004). Our study used newborn hair samples for sex hormone extraction as a novel method to reflect fetal hormone concentrations, capturing metabolic activity from the 28^th^ week of fetal development (Gareri and Koren, 2010). The fact that we were able to extract sex hormone levels for 90% of our participants renders this method feasible and promising (see Figure 1). Accordingly, this method is increasingly applied in newborn research, with empirical validation from recent studies (Deer et al., 2024; Schury et al., 2017). Apart from hair-sample analysis, hormone extraction from amniotic fluid is the only other method previously used to investigate the association between fetal hormone metabolism from the second trimester onwards and child behavior. However, as amniocentesis is mainly performed in high-risk pregnancies to diagnose fetal defects or genetic diseases, this method has clear limitations in the study of general development (Knickmeyer et al., 2005). Hormone extraction from umbilical cord blood is neither an option to capture fetal hormone levels, as it reflects perinatal concentrations of sex hormones influenced by birth (Taylor et al., 2000; van de Beek et al., 2004). Apart from the invasiveness required to access fetal body fluids, these methods’ limitation lies in their representation of only single time points of fetal hormone levels, which vary throughout the day (Bates and Herzog, 2020; Granger et al., 1999; Taylor et al., 2000). In contrast, as a non-invasive method, hair-sample analysis provides a retrospective index of long-term hormone secretion (Gao et al., 2013), offering insight into fetal endocrinological development during the third trimester.

Despite its novelty, our study has several limitations. First, the current sample size was relatively small (*n* = 63), and the educational background lacked diversity, as the sample mainly included highly educated mothers. Future research would benefit from larger and more diverse samples to strengthen the validity and applicability of the results across broader populations. Second, measuring hormones from newborn hair is an approach still in the early stages of validation with remaining challenges, such as the absence of standardized reference values, the limited amount of hair in newborns, and variability in hair growth. Note, however, that hair length and hair mass in our study did not show any statistical association with hormone concentrations (see Table S1). Moreover, steroid absorption from the amniotic fluid, influenced by fetal swallowing, urination, and steroid transfer across fetal tissues into the fetal hair matrix cannot be entirely excluded, despite washing hair samples to remove external contamination prior to analysis (Koskivuori et al., 2023). Additionally, the absence of maternal DHEA data limits interpretation. However, even though maternal influence on fetal DHEA levels via intrauterine communication cannot be ruled out entirely, fetal DHEA levels have been shown to be largely independent of maternal concentrations (Hart et al., 2023; Nieschlag et al., 1974). Another limitation of our study might be, that we used the BSID-III to assess language abilities at six months of age, which marks a very early stage in language development when language skills are only beginning to emerge, with the BSID-III largely reflecting developmental precursors. Although its validity at this age may be limited, early communicative abilities at this age, such as infant vocalizations (Werwach et al., 2021), and speech perception (Tsao et al., 2004), have been found to predict subsequent language development. Comparably, in the BSID-III, the language scale at seven months is predictive of the language scale at two years of age (Reuner and Rosenkranz, 2014), highlighting the value of early assessments. However, it remains essential to evaluate the predictive value of fetal DHEA in the course of language development, testing it at later ages.

## 5 CONCLUSIONS

The current study identifies fetal DHEA levels extracted from neonate hair samples as a relevant candidate biological marker of infant language development, with potential lasting effects for child development. Due to its anti-glucocorticoid properties, DHEA may affect fetal development by modulating cortisol activity and should be targeted in studies on prenatal programming and its impact on infant development.

## Declarations

The authors have declared no conflict of interest. This study adhered to the Declaration of Helsinki. Prior to participation, mothers provided informed consent and the experiment was approved by the ethics committee of the Paris Lodron University of Salzburg (EK-GZ: 12/2013, approved on 23^rd^ of January, 2014, Addendum: 17^th^ November, 2021). A preprint version of the manuscript is available on bioRxiv [DOI: https://doi.org/10.1101/2025.06.24.661252].

## Data availability

Non-sensitive data is available from the corresponding author upon reasonable request.

## Acknowledgements

The authors wish to thank the participating mothers, fathers, and children for their willingness to take part in this project as well as the students who helped with data collection.

