## Supplementary Information for "Fetal Dehydroepiandrosterone from Hair Samples at Birth Predicts Language Development"

**SUPPORTING INFORMATION**

### Correlation analysis of hormone concentrations with hair measures and infant age at hair sampling

S1: Spearman-Correlation Analysis of DHEA and testosterone concentrations with hair mass, hair length, and infant age at hair sampling, averaged across *m = 20* imputed data sets, *n* = 63)

| | *DHEA* | *Testosterone* |
|---|---|---|
| Hair length* | $\rho$ = .16, *p* = .23 | $\rho$ = .20, *p* = .12 |
| Hair mass | $\rho$ = .09, *p* = .46 | $\rho$ = .01, *p* = .93 |
| Infant age at hair sample collection | $\rho$ = .11, *p* = .41 | $\rho$ = -.17, *p* = .21 |

*Note.* This table presents the spearman-correlation results between each hormone level (DHEA and testosterone) and hair characteristics (mass, length), as well as age at hair sampling, averaged across *m* = 20 imputed datasets (*n* = 63). No statistically significant correlations were observed between each hormone concentration and hair mass, hair length, or age. *Analyses including hair length are based on *n* = 62 due to one missing value.

**Associations between fetal DHEA and language and cognitive skills at six months of age**

Table S2a: Multiple linear regression models examining fetal DHEA levels as main effect and interacting with infant sex on language skills at six months of age across *m* = *20* imputed datasets (*n* = 63)

| Variable | Beta | *SE* | *β* | *p* |
|---|---|---|---|---|
| Constant | 128.27 | 4.93 | - | <.001 |
| Fetal DHEA level (log-scaled) | -2.37 | 1.03 | -.28 | .02 |
| Fetal DHEA (log-scaled): Sex | -1.08 | 0.57 | -.23 | .06 |

*Note.* This table presents the pooled results from 20 separate multiple linear regression models conducted on *m* = 20 imputed datasets, examining the main effect of fetal DHEA levels and its interaction with infant sex on language skills at six months of age (*pooled* $R^2_{adj.}$ *across imputations* = .13, $p < .01$).

Table S2b: Sensitivity Analysis: Multiple linear regression models examining fetal DHEA levels as main effect and interacting with infant sex on language skills at six months of age across *m* = *20* imputed datasets (*n* = 60)

| Variable | Beta | *SE* | *β* | *p* |
|---|---|---|---|---|
| Constant | 127.85 | 4.30 | - | <.001 |
| Fetal DHEA level (log-scaled) | -2.48 | 0.90 | -.35 | <.01 |
| Fetal DHEA (log-scaled): Sex | -0.84 | 0.50 | -.21 | .10 |

*Note.* Of *n* = 63 participants, *n* = 3 participants were identified as statistical outliers (SD > 2.5) and excluded from additional sensitivity analysis. Pooled results of multiple linear regression models on 20 imputed datasets (*pooled* $R^2_{adj.}$ *across imputations* = .16, $p < .01$).

Table S3: Multiple linear regression models examining fetal DHEA levels as main effect and interacting with infant sex on cognitive skills at six months of age across *m = 20* imputed datasets ($n$ = 63)

| Variable | Beta | *SE* | $\beta$ | $p$ |
|---|---|---|---|---|
| Constant | 110.18 | 6.63 | - | <.001 |
| Fetal DHEA level (log-scaled) | 1.20 | 1.39 | .11 | .39 |
| Fetal DHEA (log-scaled): Sex | -0.72 | 0.78 | -.12 | .36 |

*Note.* This table presents the pooled results from 20 separate multiple linear regression models conducted on $m$ = 20 imputed datasets, examining the main effect of fetal DHEA levels and its interaction with infant sex on cognitive skills at six months of age (*pooled* $R^2_{adj.}$ *across imputations* = -.01, $p$ = .50). As no cases with standardized residuals beyond *2.5 SD* were identified in any of the imputed regression models, sensitivity analysis was not warranted.

**Associations between fetal testosterone and language skills at six months of age**

Table S4a: Multiple linear regression models examining fetal testosterone levels as main effect and interacting with infant sex on language skills at six months of age across *m = 20* imputed datasets (*n* = 63)

| Variable | Beta | *SE* | *β* | *p* |
|---|---|---|---|---|
| Constant | 113.69 | 2.24 | - | <.001 |
| Fetal testosterone level (log-scaled) | 1.13 | 1.08 | .16 | .30 |
| Fetal testosterone (log-scaled): Sex | -1.08 | 1.26 | -.13 | .40 |

*Note.* This table presents the pooled results from 20 separate multiple linear regression models conducted on *m* = 20 imputed datasets, examining the main effect of fetal testosterone levels and its interaction with infant sex on language skills at six months of age (*pooled* $R^2_{adj.}$ *across imputations* = -.01, *p* = .52).

Table S4b: Sensitivity Analysis: Multiple linear regression models examining fetal DHEA levels as main effect and interacting with infant sex on language skills at six months of age across *m = 20* imputed datasets (*n* = 60)

| Variable | Beta | *SE* | *β* | *p* |
|---|---|---|---|---|
| Constant | 113.56 | 1.95 | - | <.001 |
| Fetal testosterone level (log scaled) | 1.17 | 0.94 | .19 | .22 |
| Fetal testosterone (log scaled): Sex | -1.62 | 1.13 | -.22 | .16 |

*Note.* Of *n* = 63 participants, *n* = 3 participants were identified as statistical outliers (SD > 2.5) and excluded from additional sensitivity analysis. The pooled results of multiple linear regression models on 20 imputed datasets are presented (*pooled* $R^2_{adj.}$ *across imputations* = .01, *p* = .29).
